# Tensorial blind source separation for improved analysis of multi-omic data

**DOI:** 10.1101/300277

**Authors:** Andrew E Teschendorff, Jing Han, Dirk S Paul, Joni Virta, Klaus Nordhausen

## Abstract

There is an increased need for integrative analyses of multi-omic data. Although several algorithms for analysing multi-omic data exist, no study has yet performed a detailed comparison of these methods in biologically relevant contexts. Here we benchmark a novel tensorial independent component analysis (tICA) algorithm against current state-of-the-art methods. Using simulated and real multi-omic data, we find that tICA outperforms established methods in identifying biological sources of data variation at a significantly reduced computational cost. Using two independent multi cell-type EWAS, we further demonstrate how tICA can identify, in the absence of genotype information, mQTLs at a higher sensitivity than competing multi-way algorithms. We validate mQTLs found with tICA in an independent set, and demonstrate that approximately 75% of mQTLs are independent of blood cell subtype. In an application to multi-omic cancer data, tICA identifies many gene modules whose expression variation across tumors is driven by copy number or DNA methylation changes, but whose deregulation relative to the normal state is independent such alterations, an important finding that we confirm by direct analysis of individual data types. In summary, tICA is a powerful novel algorithm for decomposing multi-omic data, which will be of great value to the research community.

## Background

Omic data is now most often generated in a multi-dimensional context. For instance, for the same individual and tissue-type one may measure different data-modalities (e.g. genotype, mutations, DNA methylation or gene expression), which may help pinpoint disease driver genes [1]. Alternatively, for the same individual, the same data-type may be measured across different tissues or cell-types [2, 3], which may help identify the most relevant cell-types or tissues for understanding disease aetiology. We refer to all of these types of multi-dimensional data generally as “multi-way” or “multi-omic” data, and when samples and molecular features are matched, the data can brought into the form of a multi-dimensional array, formally known as a tensor [4].

While several statistical algorithms for the analysis of multi-way or tensorial data are available [5, 6, 4, 7], their application to real data has been challenging. There are mainly three reasons for this. First, the associated multi-way datasets are often very large and how well the algorithms perform on such large sets is currently still unclear. Second, the algorithms can be computationally demanding, compromising their benefit-to-cost ratio [4]. Third, interpreting the output of these algorithms requires an in-depth understanding of the underlying methods. Exacerbating this problem, most available software packages are not user-friendly, requiring the user to possess such in-depth understanding in order to extract the relevant biological information. Beyond these technical challenges, there is also a lack of comparative studies, making it difficult to choose the appropriate algorithm for the task in question.

To help address some of these outstanding challenges, we here consider and evaluate a novel data tensor decomposition algorithm [8, 9], which is based on the framework of blind source separation (BSS), and specifically that of Independent Component Analysis (ICA) [10]. Although common BSS techniques such as non-negative matrix factorization (NMF) and ICA have been successfully applied to a wide range of single omic data types, including e.g. gene expression [11, 12, 13, 14, 15, 16], DNA methylation [17] and mutational data [18], their application to multi-way data is largely unexplored [19]. In the case of single omic datasets, the improved performance of ICA over non-BSS techniques like PCA owes primarily to the non-Gaussian and often sparse nature of biological sources of variation, which means that statistical deconvolution of biological samples benefits from non-linear decorrelation measures such as statistical independence (as used in ICA) [13]. It is therefore natural to consider analogous ICA algorithms for multi-way data, as we do here, since these may also lead to improved inference.

In order to assess this, we here benchmark our novel tensorial BSS algorithm against some of the most popular and powerful algorithms for inferring sources of variation from multi-omic data, including JIVE (Joint and Individual Variation Explained) [5], PARAFAC (Parallel Factor Analysis) [6, 4], iCluster [7] and Canonical Correlation Analysis (CCA) [20, 21, 22]. Each of these algorithms has particular strengths and weaknesses, which render comparisons between them highly non-trivial. For instance, a limitation of CCA is that it can only infer common sources of variation between data-types or tissues, in contrast to JIVE or PARAFAC which can infer both joint as well as individual sources of variation. On the other hand, JIVE and CCA can be run on multiple data matrices with different numbers of molecular features, while PARAFAC and iCluster require matched sets of features (and samples) for each data-type. Model complexity also differs substantially between methods, with PARAFAC exhibiting a much lower model complexity than an algorithm such as iCluster. Thus, a comparison of all of these methods is of paramount interest, and here we do so in a tensorial context, i.e. one where the multi-way data is defined over a matched set of molecular features (e.g. genes or CpGs) and samples across all data-types, allowing the data to be brought into the form of a tensor. Specifically, we shall here consider order-3 data tensors, i.e. data which can be brought into the form of an array with 3 dimensions (often called “modes”). In our evaluation and comparison of all multi-way algorithms, we consider both simulated data as well as data from real epigenome-wide association studies (EWAS). We further illustrate potential uses of our tensorial BSS algorithm for (i) the detection of cell-type independent and cell-type specific methylation quantitative trait loci (mQTLs) in multi cell-type or multi-tissue EWAS, and (ii) the detection of cancer gene modules deregulated by copy-number and DNA methylation changes.

## Results

### Tensorial ICA outperforms JIVE, PARAFAC, iCluster and CCA on simulated data

Tensorial ICA (tICA) aims to infer from a data-tensor, statistically independent sources of data variation, which should better correspond to underlying biological factors (**Methods**). Indeed, since biological sources of data variation are generally non-Gaussian and often sparse, the statistical independence assumption implicit in the ICA formalism can help improve the deconvolution of complex mixtures and thus better identify the true sources of data variation (**Fig.1**). As with ordinary ICA itself, there are different ways of implementing tICA, and we here consider two different flavours, called tensorial fourth-order blind identification (tFOBI) and tensorial joint approximate diagonalization of high-order eigenmatrices (tJADE) (**Methods**). Specifically, we here consider two modified versions of these, whereby tensorial PCA is applied as a noise reduction step (also called whitening) prior to implementing tICA, resulting in two algorithms we call tWFOBI and tWJADE (**Methods**).

**Figure 1.**
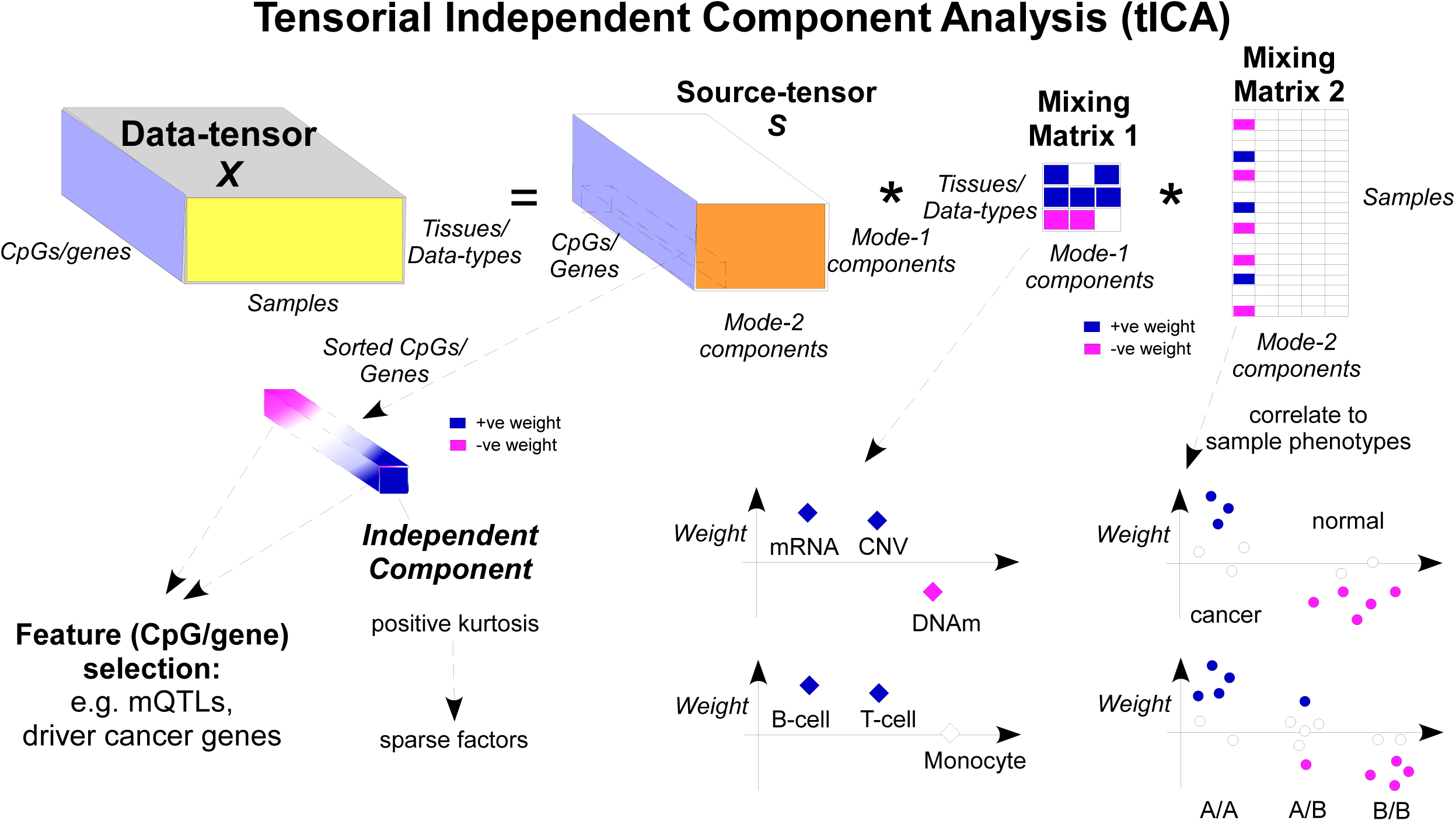
Decomposing data tensors using Independent Component Analysis. Tensorial ICA (tICA) works by decomposing a data-tensor, here depicted as an order-3 tensor with 3 dimensions representing features (CpGs/genes), samples and tissue or data-type, into a source tensor S and two mixing matrices defined over tissue/data-type and samples, respectively. The key property of tICA is that the independent components in S are as statistically independent from each other as possible. Statistical independence is a stronger criterion than linear decorrelation and allows improved inference of sparse sources of data variation. Positive kurtosis can be used to rank independent components to select the most sparse factors. The largest absolute weights within each independent component can be used for feature selection, while the corresponding component in the mixing matrices inform about the pattern of variation of this component across tissue/data-types and samples, respectively. In the latter case, the weights can be correlated to sample phenotypes such as normal/cancer status or genotype. For the first mixing matrix, the weights inform us about the relation between data-types (e.g. if the copy-number change is positively correlated with gene expression), or in the case of a multi-cell EWAS, whether mQTLs are cell-type independent or not. Abbreviations: +ve=positive, -ve=negative.

First, we tested the two tICA algorithms, as well as tPCA, on simulated multi-way data consisting of two different data matrices defined over the same 1000 features (genes) and 100 samples (**Methods**). The data for the 2 matrices was generated with a total of 4 sources of variation, 2 for each matrix, and with 1 source in each data matrix describing joint variation, driven by a total of 100 genes. A total of nine different noise levels were simulated, ranging from a high signal-to-noise ratio (SNR) regime (SNR=3, NoiseLevel=1) to a low SNR regime (SNR=0.6, Noise-Level=5). For each noise level, a total of 1000 Monte-Carlo runs were performed. In each run, we compared the multi-way algorithms in terms of their sensitivity (SE) and specificity (SP) to detect the 50 genes driving the joint variation. We did not consider the corresponding performance measures for the individual variation (i.e. the variation specific to one data-type), because not all algorithms infer sources of individual variation (e.g. CCA), thus precluding direct comparison between them, and because identifying sources of joint variation is always the main purpose of multi-way algorithms. The number of components chosen for each method and the number of genes selected within components to compute SE and SP is explained in detail in **Methods**. SE and SP values for joint variation of each algorithm and noise-level were averaged over the 1000 runs (**Methods**). Benchmarking tICA and tPCA against PARAFAC, CCA, JIVE and iCluster, we observed that for low noise levels all algorithms performed similarly, except PARAFAC which exhibited significantly worse SE and SP values (**Fig.2A,2C**). For larger noise levels, we observed worse performance for JIVE, CCA and iCluster compared to the two different tICA methods (tWFOBI & tWJADE) (**Fig.2, Methods**). Differences in SE and SP between the tICA methods and JIVE, CCA, iCluster and PARAFAC were statistically significant (**Fig.2B,2D**). On this data, and since tICA uses tensorial PCA (tPCA) as a preliminary step, we did not observe substantial difference between tPCA and tICA (**Fig.2**). We note that in this evaluation on the simulated data we did not consider sparse-CCA (SCCA), since the sparsity itself does not optimize sensitivity and thus sCCA would perform substantially worse than CCA (data not shown). Results were unchanged if we replaced Gaussian distributions (as the sources of variation) with supergaussian Laplace distributions, indicating that results are not dependent on the type of data distribution (**fig.S1 in SI**).

**Figure 2.**
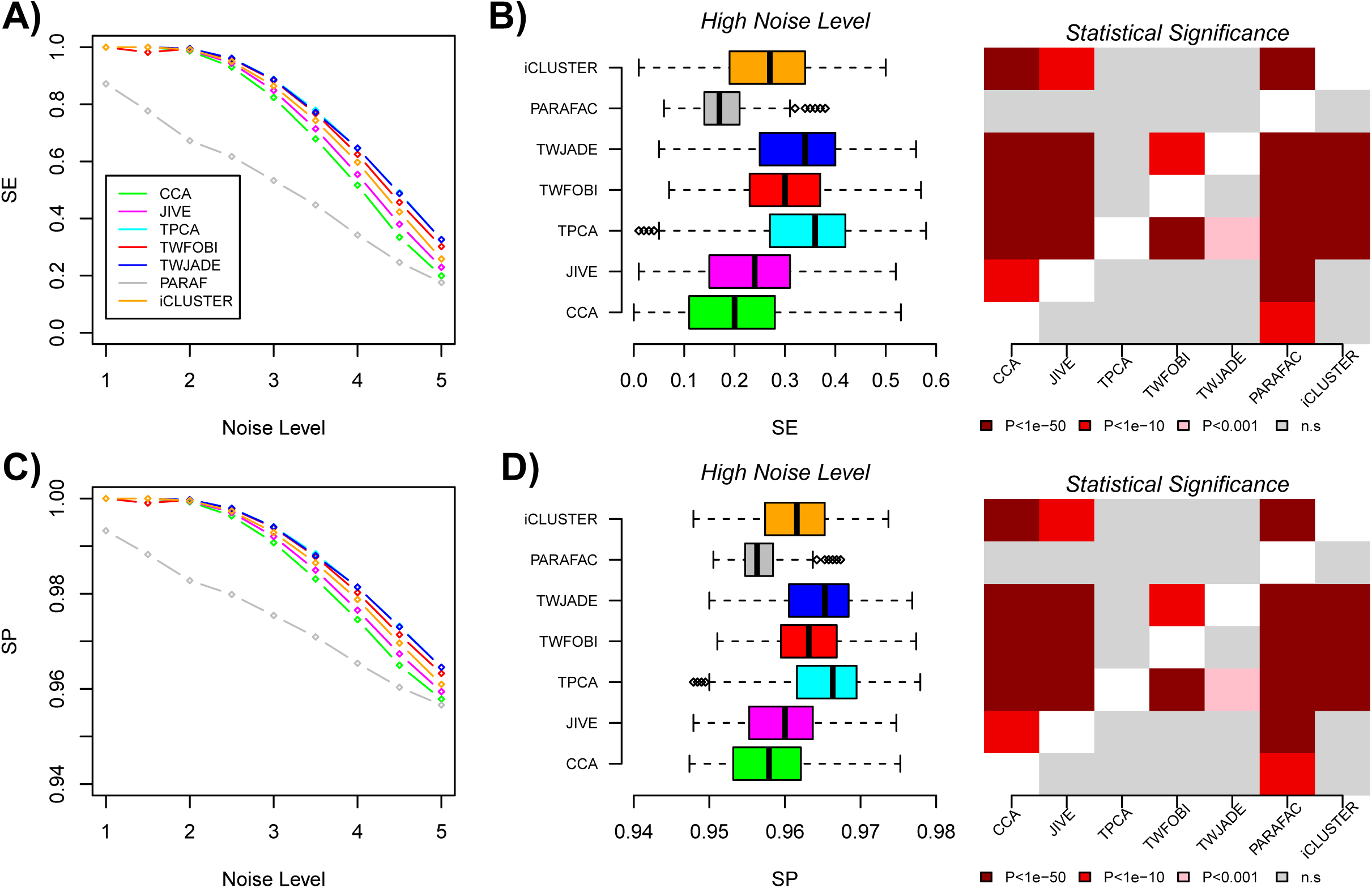
Comparison of multi-way algorithms on simulated data. **A**) Sensitivity (SE) versus noise level (x-axis) for 7 different methods as indicated, as evaluated on simulated data (data points are averages over 1000 Monte-Carlo runs). In each case the data-tensor was of size 2 × 100 × 1000, i.e. 2 data-types, 100 samples and 1000 genes. **B) Left panel:** Boxplots of SE values for the same 7 methods for the largest noise level(=5). Each box contains the SE-values over the 1000 Monte-Carlo runs. **Right panel:** Corresponding heatmap of P-values of significance for each pairwise comparison of methods. P-values computed from a one-tailed Wilcoxon rank sum test. For each entry specified by a given row and column, the alternative hypothesis is that the method specified in the row has a higher SE than the method specified in the column. **C-D** As A-B), but for the specificity (SP).

### Tensorial PCA/ICA reduces running time over JIVE, PARAFAC and iCluster

Using the same simulated data, we further compared the algorithms in terms of their running times. A detailed comparison is cumbersome because the parameters specifying the number of components to search for are not directly comparable and differ substantially between methods. Nevertheless, using reasonable parameter choices for the simulated model above, we found that tPCA and tICA substantially speed up inference over methods such as JIVE or iCluster (**Table 1**). In fact, even when specifying a larger number of components for tPCA/tICA, compared to PARAFAC, JIVE or iCluster, the latter were substantially slower (**Table 1**), whilst also exhibiting marginally worse SE and SP values (**Fig.2**). In general, we observed tICA methods to be at least 50 times faster than PARAFAC, and at least 100 times faster than JIVE and iCluster (**Table 1**). For much larger datasets, we found application of iCluster to be computationally demanding and not practical. Thus, in subsequent analyses on real datasets we decided to benchmark tPCA/tICA against PARAFAC, CCA, SCCA and JIVE.

**Table 1.**
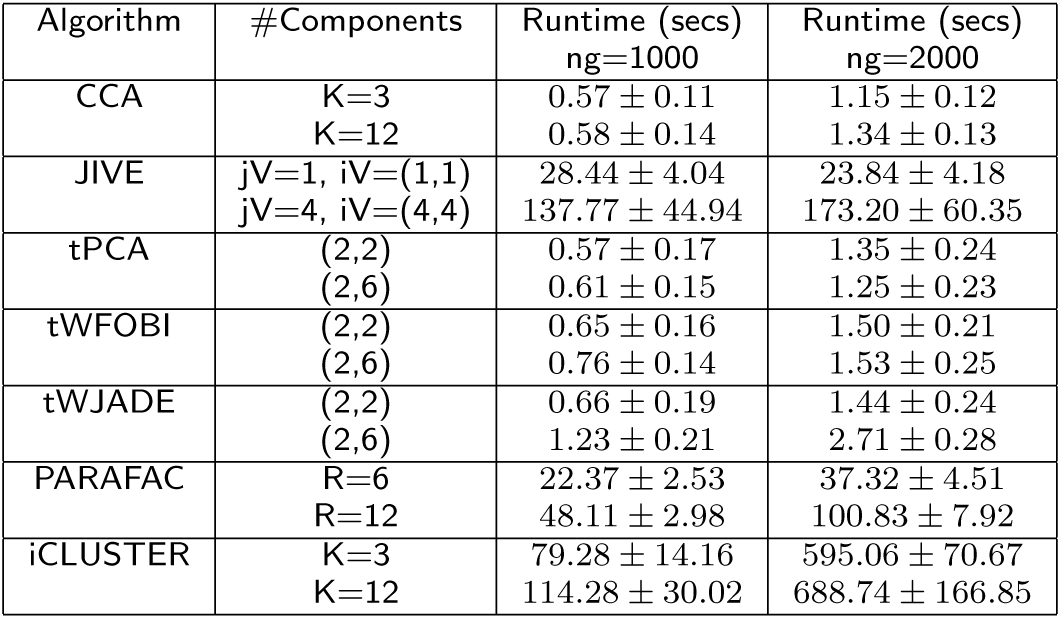
Comparison of running times of multi-way algorithms. Table compares 7 multi-way algorithms in terms of the running times to infer components of variation (RunTime) in the simulation model considered in Fig.2. Estimates are medians and median absolute deviations over 100 Monte-Carlo runs for the case where the signal-to-noise ratio is 1 (i.e. NoiseLevel==3 in Fig.2). Second column specifies the parameter values for the number of components used in each algorithm. In the case of the first rows for each method: for CCA, 3 sets of canonical vector pairs (K=3), for JIVE rank of joint variation (jV)=1, rank of individual variation (iV) for each data type=1, for TPCA, TWFOBI and TWJADE we inferred 2 and 2 components in the data-type and sample dimensions respectively, for PARAFAC the rank of decomposition was R=6 and for iCLUSTER the maximum number of clusters K was set to 3. For the second rows, total number of components are exactly matched (12) for all methods. Running times are reported for two scenarios differing in the number of genes ng, as indicated, and were obtained on a Dell PowerEdge R830 with Intel Xeon E5-4660 v4 2.2GHz processors.

### Tensorial ICA exhibits improved power in a real multi-tissue smoking EWAS

Next, we asked if tPCA/tICA also leads to improved power on real data. Objective evaluation on real data is challenging due to the difficulty of defining a gold-standard set of true positive associations. Fortunately however, a meta-analysis of several smoking EWAS in blood has demonstrated that smoking-associated DMCs (smkDMCs) are highly reproducible, defining a gold-standard set of 62 smkDMCs (**Methods**) [23]. Recently, we also showed that effectively all 62 smkDMCs are associated with smoking exposure if DNAm is measured in buccal samples [2]. Thus, one way to objectively compare algorithms is in terms of their sensitivity to identify these 62 smkDMCs in a matched blood-buccal EWAS consisting of Illumina 450k DNAm profiles for a total of 152 women (**Methods**, [2]). Because there are two distinct samples (1 blood + 1 buccal) per individual, most of the variation is genetic. Hence, to reduce this background genetic variation, we first computed the SE-values on a reduced data matrix obtained by combining the 62 smkDMCs with 1000 randomly selected non-smoking associated CpGs (a total of 100 Monte-Carlo randomizations). We considered both the maximum SE value attained by a component, as well as the overall SE obtained by combining selected CpGs from components significantly enriched for smkDMCs (**Methods**). This revealed that JIVE, CCA/SCCA and PARAFAC were all superceded by tPCA and tICA (**Fig.3A-B**). Differences between tPCA and tICA were generally not significant (**Fig.3A**), although tWFOBI attained higher combined SE values than tPCA and tWJADE (**Fig.3B**).

**Figure 3.**
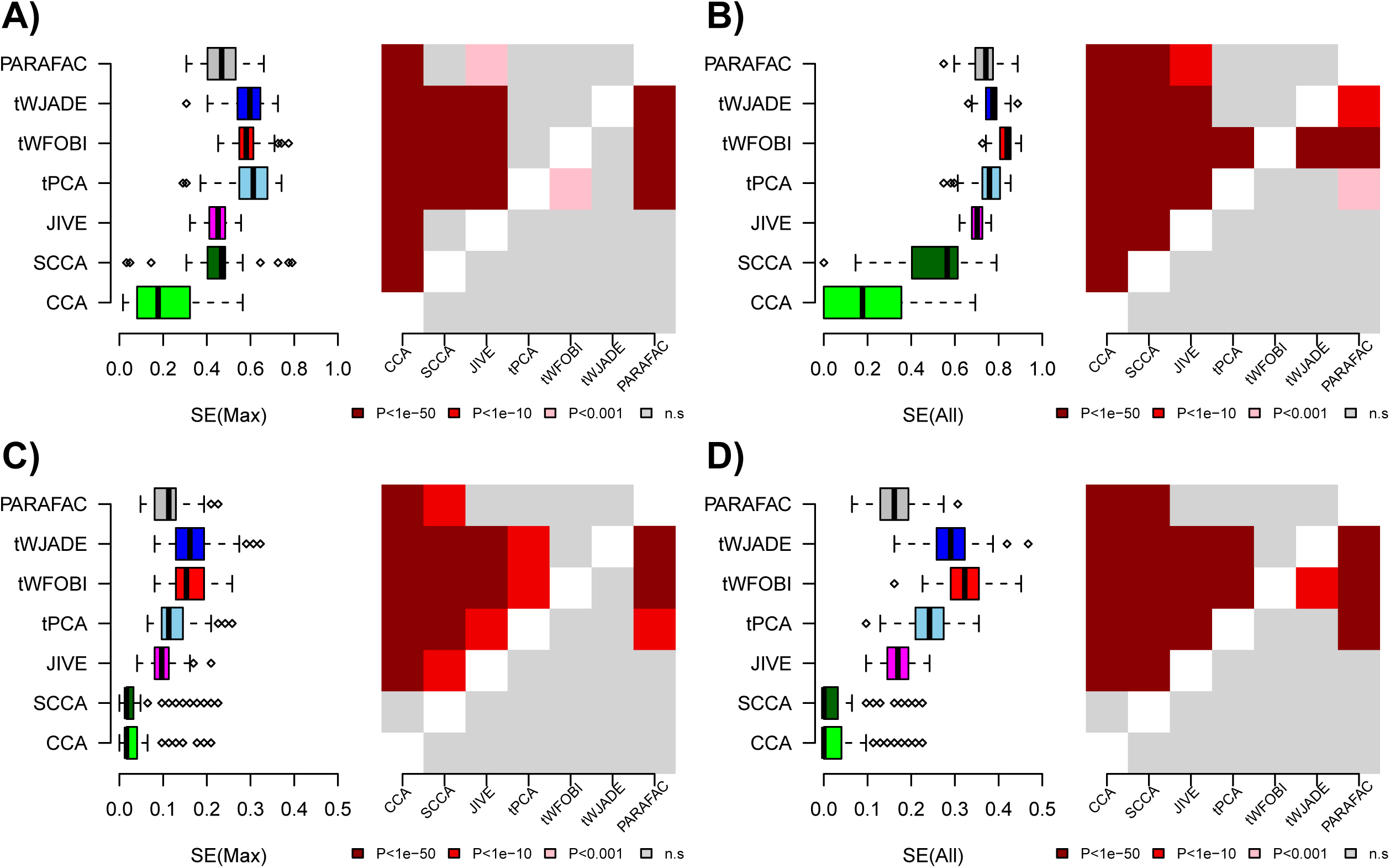
Comparison of multi-way algorithms on a multi-tissue smoking EWAS. **A)** Boxplot of sensitivity (SE) values for each of the 7 methods as applied to the data tensors of dimension 2 × 152 × 1062 (2 tissues, 152 samples, 1000 randomly selected non-smkDMCs + 62 smkDMCs) and for 100 different selections of non-smkDMCs. SE(Max) denotes that maximum sensitivity to capture 62 smkDMCs among all inferred components. Right panel is a heatmap of the corresponding one-tailed paired Wilcoxon rank sum test, benchmarking the SE values of each method (y-axis) against each other method (x-axis). **B)** As A), but now for the combined sensitivity (SE(All)) obtained from all enriched components. **C-D)** As A-B), but now for data-tensors of dimension 2 × 152 × 10062 and for 100 randomly selected 10000 non-smkDMCs.

Next, we scaled up the data matrices by combining the 62 smkDMCs with a larger set of 10000 non-smkDMCs, recomputing the SEs (again for 100 different Monte-Carlo selections of 10000 non-smkDMCs). As expected, with an increase in the number of CpGs, the SE of all algorithms dropped, likely driven by increased confounding due to genetic variation (**Fig.3C-D**). With the increase in probe number, tICA (tWFOBI & tWJADE) outperformed not only JIVE, PARAFAC and CCA/SCCA, but also tPCA (**Fig.3C-D**), in line with the increased sparsity of the smoking-associated source of variation.

In order to illustrate how the output produced by tICA can be used for valuable inference, we focus on a particular Monte-Carlo run and a specific component (estimated using tWJADE), which obtained a high sensitivity for smkDMCs (component-12, **Fig.4A**). We note that the two ICs *S*_1,12,*i*_ and *S*_2,12,*i*_ exhibited a less correlative structure than the corresponding components projected onto the blood and buccal dimensions, demonstrating that tWJADE does indeed identify components that are less statistically dependent (**Fig.4A**). Confirming the high sensitivity of these ICs, the 62 smkDMCs were highly enriched among CpGs with the largest absolute weights in any one of the two ICs (**Fig.4A**, Fisher-test *P* < 1*e* −36, SE=41/62~0.66). We further verified that the 41 enriched smkDMCs exhibited strong Pearson correlations between their DNAm profiles in blood and buccal, as required since smoking exposure is associated with similar DNAm patterns in these two tissue types (**Fig.4B**, [2]). Further confirming that component-12 is associated with smoking exposure, we correlated the weights of the corresponding column of the estimated mixing matrix with two different measures of smoking exposure, demonstrating in both cases a strong association (**Fig.4C**). Thus, application of tICA on DNAm data results in components that are readily interpretable in terms of their associations with known smoking exposures across features and samples.

**Figure 4.**
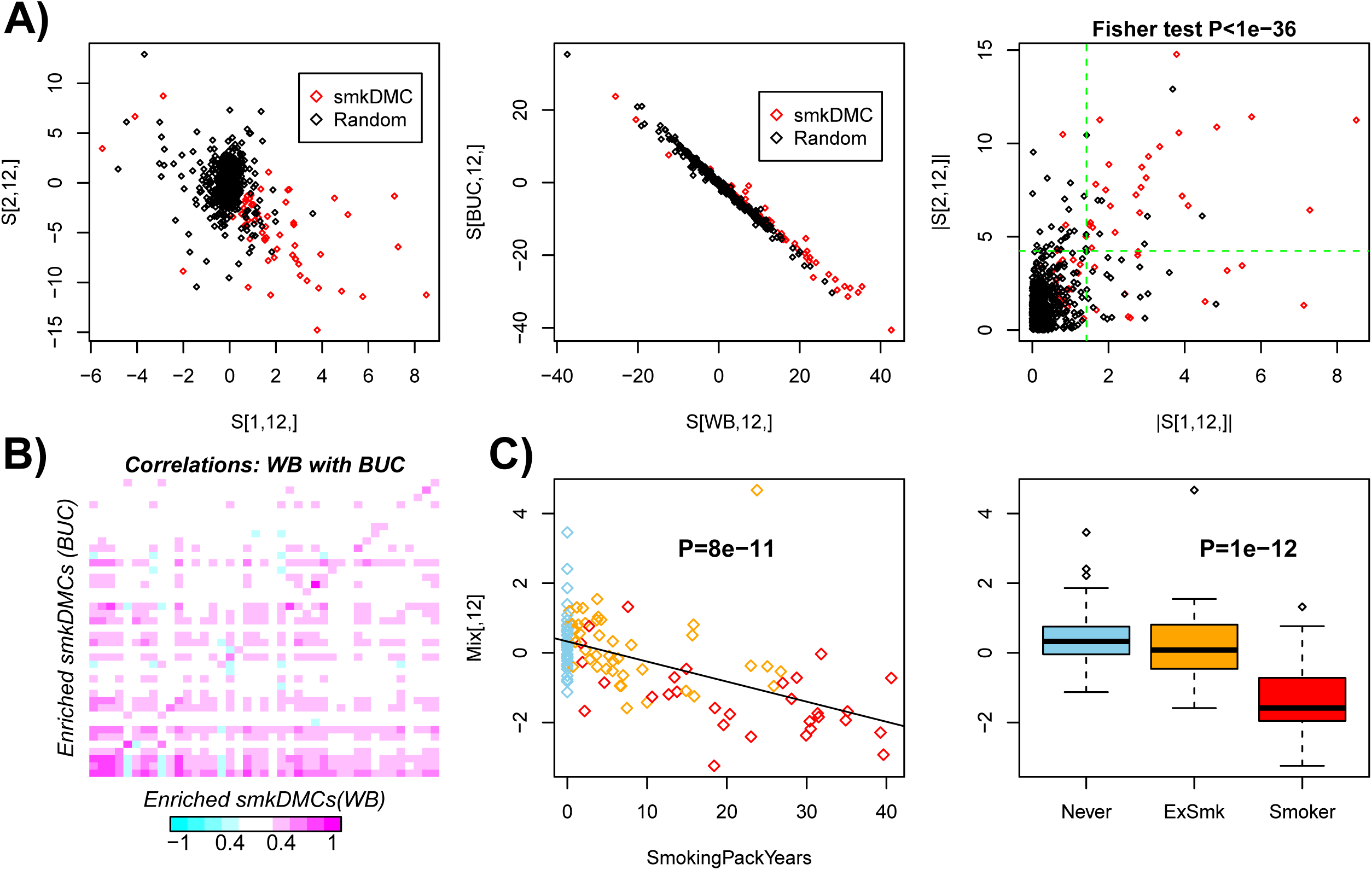
Validation of tensorial ICA on multi-tissue smoking EWAS. **A)** Left panel: Scatterplot of the weights of an estimated independent components *S*_1,12,_*_i_* and *S*_2,12,_*_i_* from the data tensor of dimension 2 × 152 × 1062, with mode-1 representing tissue-type, mode-2 the different women and mode-3 representing the CpGs. Red denotes the smkDMCs. Middle panel: As left panel, but now for the rotated tensor, projecting the data onto the whole blood and buccal dimensions, demonstrating the strong correlation between the DNAm variation in whole blood (WB) and buccal (BUC) tissue. Right panel: As left panel, but now for the absolute weights, and with green dashed lines representing the cutoff point selecting the 62 CpGs with the largest absolute weights. There are in total 41 smkDMCs among the 3 larger quadrants, corresponding to a sensitivity of 41/62=0.66, with the enrichment P-value given above the plot. **B)** Pearson correlation heatmap of the 41 smkDMCs between whole blood (WB) and buccal (BUC) tissue, with correlations computed over the 152 samples. **C)** Plots of the 12th IC of the mixing matrix in sample-space (y-axis) against smoking exposure for the 152 samples. Left panel is for smoking-pack-years, right panel is for smoking status: never smokers, ex-smokers and smokers at sample draw. P-values are from linear regressions.

### tICA identifies mQTLs in a multi cell-type EWAS

Having established the better performance of tICA over other state-of-the-art methods, we next considered the application of tICA (specifically tWFOBI) in an EWAS of 47 healthy individuals, for which 3 purified cell-types (B-cells, T-cells and Monocytes) had been profiled with Illumina 450k DNA methylation (DNAm) beadarrays [3] (**Methods**). We chose tWFOBI over tWJADE because of its computational efficiency (**Table 1**). Given that 3 cell-types were measured for each individual, the expectation is that a significant amount of inter-individual variation in DNAm would correlate with genetic variants (i.e. methylation quantitative trait loci-mQTLs) [24]. Thus, it is important to evaluate the ability of tICA to detect mQTLs and to determine whether these are blood cell-subtype specific or not. Applying tWFOBI to the 3 cell-type × 47 sample × 388618 probe data-tensor, we inferred a total of 11 ICs in sample-mode space (yielding 33 ICs across sample and cell-type modes combined). For each of these 11 ICs in each cell-type, we ranked probes according to their absolute weights and tested enrichment of the top-500 probes against a high-quality list of 22,245 mQTLs as derived in [25] (**Methods**). This high-confidence list of mQTLs all passed a very stringent unadjusted P-value threshold of P=1e-14 in each of five different human cohorts, encompassing five different age-groups [25]. We observed strong statistical enrichment for mQTLs in many ICs (**Fig.5A**). We also tested separately for enrichment of chromosomes. This revealed enrichment, notably of chromosomes 6 and 21, but also of 1, 4, 7 and 8 (**Fig.5B**). For instance, IC-9 was enriched for mQTLs and chromosome-1 in all 3 cell-types (**Fig.5A-B**). Supporting this, we found a clear example of a cell-type independent mQTL mapping to the 1q32 locus of the *PM20D1 gen*e (**Fig.5C**), a major GWAS locus associated with Parkinson’s disease [26]. Focusing on chromosome-6, another cell-type independent mQTL mapped to *MDGA1* (**fig.S2 in SI**), a major susceptibility locus for schizophrenia [27]. Other mQTLs driving ICs were cell-type specific, e.g. mQTLs mapping to *ATXN1* and *SYNJ2* were dominant in the independent components projected along B-cells, but not among T-cells or Monocytes (**fig.S3 in SI**). Although assessing whether mQTLs are truly cell-type independent or cell-type specific is not possible without genotype information, we nevertheless estimated, based on the IC weight distribution of the mQTLs across cell-types, that approximately 75% of the mQTLs enriched in ICs were cell-type independent (**fig.S4 in SI**). This estimate of the non-specificity of blood cell subtype mQTLs is similar to the one obtained by a previous study (≥ 79%) using neutrophils, monocytes and T-cells [28].

**Figure 5.**
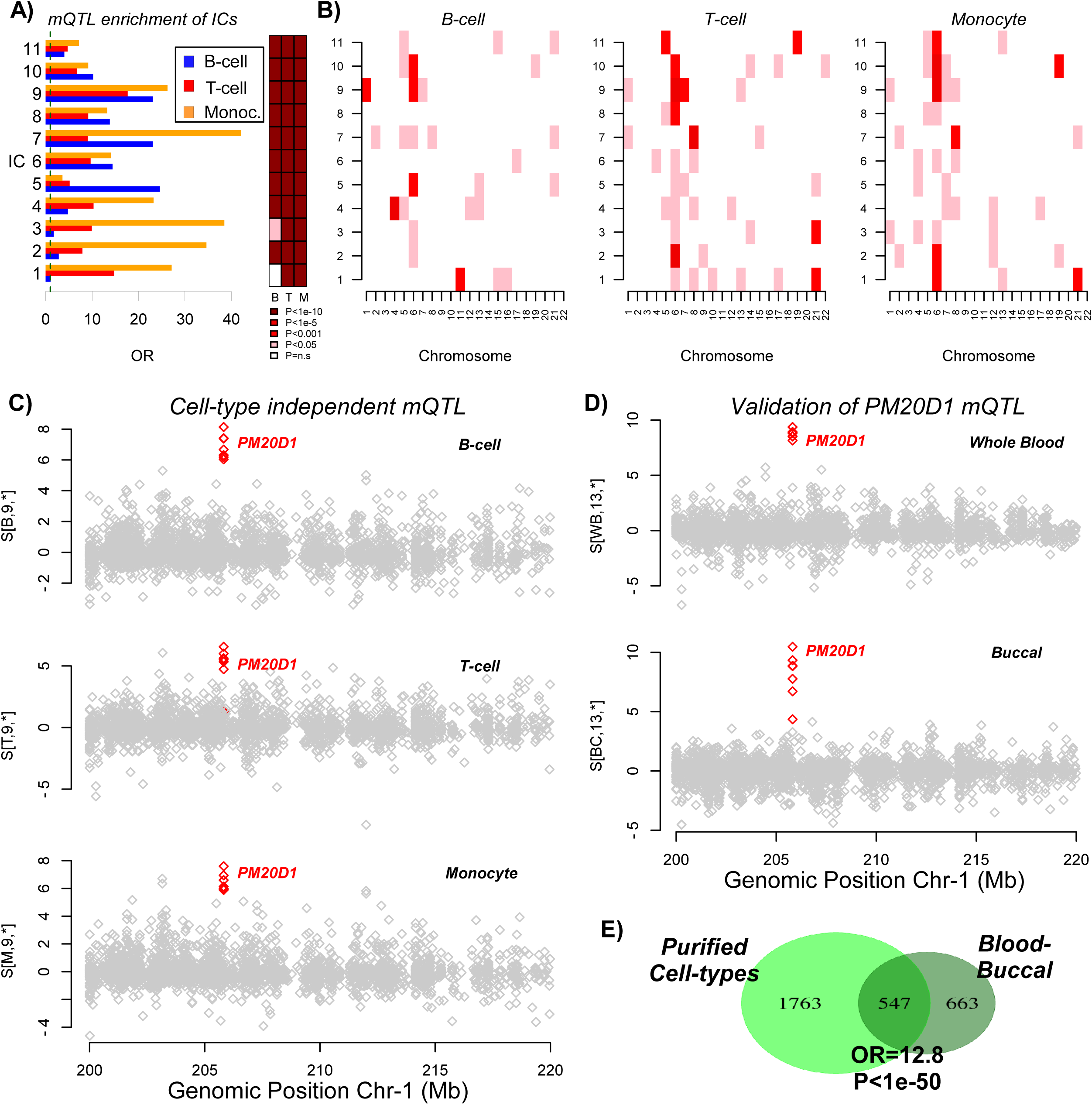
Tensorial ICA identifies components enriched for mQTLs in an EWAS of purified cell-types. **A)** Barplots of the odds ratio (OR) of enrichment of the top-ranked 500 CpGs for mQTLs in each of the 11 ICs and cell-types, as indicated. Right panel shows a corresponding heatmap indicating the P-values of enrichment as estimated using a one-tailed Fisher’s exact test. **B)** Heatmaps of enrichment P-values of the top-ranked 500 CpGs from each IC for chromosomes. Significance of P-values is indicated in different colors using same scheme as in A). **C)** An example of a cell-type independent mQTL mapping to chromosome-1. Plots show the weights in the corresponding components for B-cell, T-cell and Monocyte, respectively, with the selected CpGs mapping to the mQTL indicated in red. **D)** Validation of the mQTL in C) in an independent blood-buccal EWAS. **F)** Venn Diagram showing the overlap of mQTLs derived from the ICs in the purified cell-type EWAS with those derived from the blood-buccal EWAS. Odds ratio (OR) and one-tailed Fisher-test P-value of overlap are given.

Next, we decided to validate the found mQTLs using an independent dataset. To this end, we applied tWFOBI to the blood-buccal EWAS considered earlier. We inferred a source-tensor of dimension 2×26×447259, i.e. a total of 52 ICs, defined over 2 tissue-types and 26 components in sample-mode space. As before, we observed very strong enrichment, notably for the same chromosomes 6 and 21 (**fig.S5 in SI**). The previously found mQTL at the *PM20D1* locus, was also prominent in one of the inferred ICs in this blood-buccal EWAS, confirming its validity and further supporting that this mQTL is cell-type independent (**Fig.5D**). Overall, from the pure blood cell subtype EWAS we detected a total of 1763 mQTLs, of which 547 were also observed in the blood-buccal EWAS (OR=12.8, Fisher-test *P* < 1*e* − 50, **Fig.5E**). Thus, we can conclude that tWFOBI is able to identify components of variation across cell-types and samples that capture a significant number of mQTLs, without the need for matched genotype information.

### tICA outperforms JIVE and PARAFAC in their sensitivity to detect mQTLs

Given the ability of tICA to detect mQTLs, we next benchmarked the performance of all algorithms in terms of their sensitivity to detect mQTLs in the EWAS of three purified blood cell subtypes considered earlier. Because of the presence of three cell-types, for this analysis we excluded CCA and sCCA since these methods are designed for the case of only two data-matrices. As before, we computed two sensitivity measures to detect the 22245 mQTLs from the Aries database [25], designed to assess the overall sensitivity across all inferred components, and another designed to assess the maximum sensitivity attained by any single component. Varying the number of top ranked selected CpGs in components from 500 up to 22245, we observed that over the whole range tFOBI and tJADE were optimal, clearly outperforming both PARAFAC and JIVE (**Fig.6A**). The maximum sensitivity attained by any individual component was also best for the tICA methods (**Fig.6B**). To better evaluate the enrichment of these components for mQTLs, we also considered the ratio of the sensitivity to the maximum possible sensitivity, recording the maximum value attained by any component. This demonstrated that for the case of selecting the top-500 CpGs, that components inferred using tICA could capture over 60% of the maximum possible number of mQTLs, i.e over 60% of the 500 CpGs mapped to mQTLs (**Fig.6C**). In contrast, JIVE components only contained just over 40% of mQTLs (**Fig.6C**). We note that although the performance of JIVE could be significantly improved by also including the components of individual variation, that approximately 80% of mQTLs have been estimated to be independent of blood cell subtype [28], supporting the view that JIVE is less sensitive to capture cell-type independent mQTLs. All these results were stable to repeated runs of the algorithms, as only PARAFAC exhibited variation between runs, yet this variation was relatively small (**fig.S6 in SI**).

**Figure 6.**
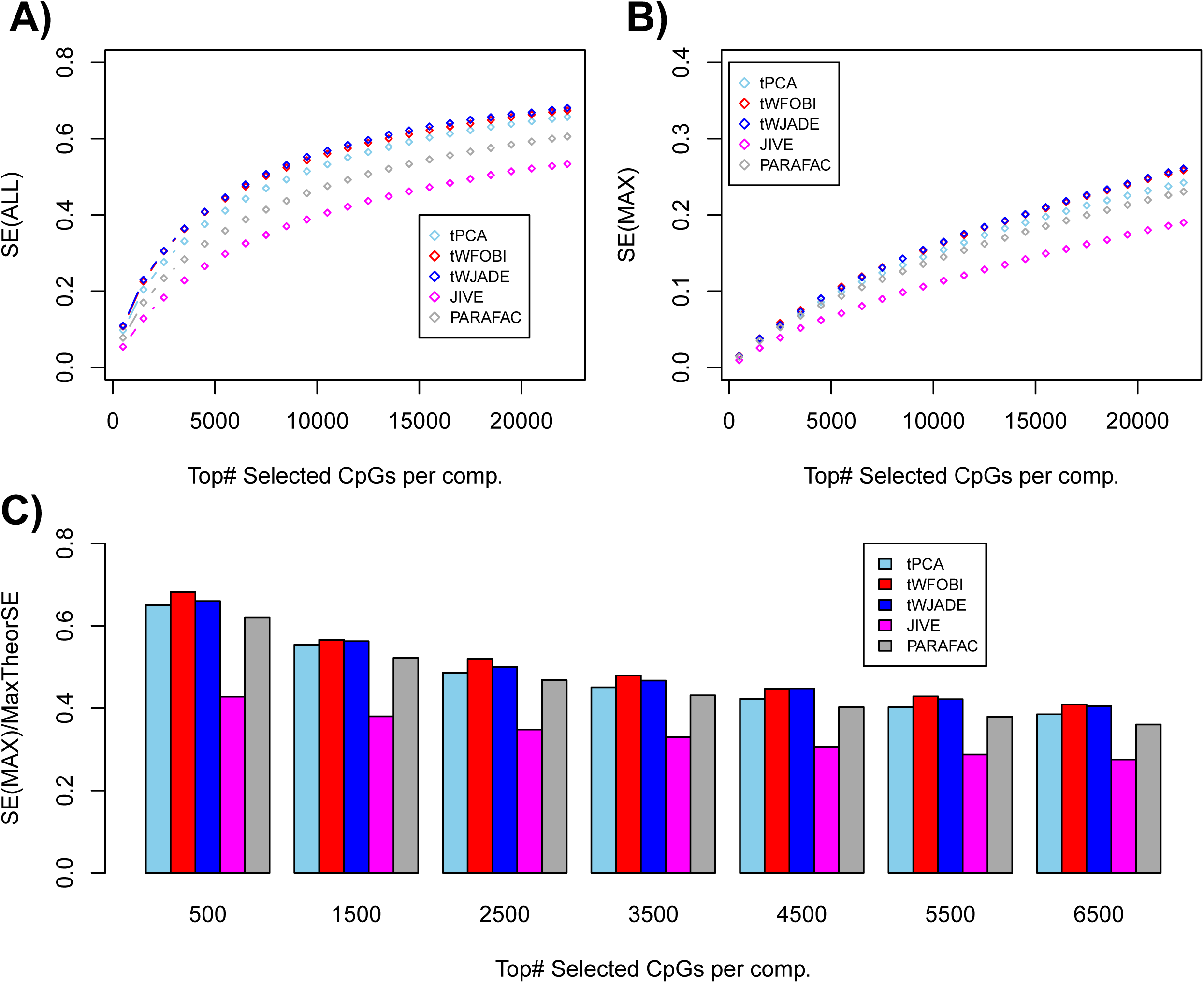
tICA outperforms JIVE and PARAFAC in detecting mQTLs. **A)** Plot of the overall sensitivity (SE(ALL),y-axis) against the number of top ranked CpGs selected in a component (x-axis) for 5 different algorithms. **B)** As A) but now for the maximum sensitivity attained by any single component (SE(MAX),y-axis). **C)** Barplot of the maximum sensitivity attained by any single component expressed as a fraction of maximum possible value given the number of selected top-ranked CpGs per component.

Next, we repeated the same sensitivity analysis to detect mQTLs in our buccal-blood EWAS, now also including CCA and sCCA (as there are only 2 tissue/cell types). Confirming the previous analysis, tICA methods outperformed JIVE and PARAFAC by over 20% in terms of the overall sensitivity, whilst also attaining a better sensitivity at the individual component level (**fig.S7 in SI**). Of note, the sensitivity of both CCA and sCCA was substantially worse, due mainly to only the top canonical vector being significant.

### Application of tICA to multi-omic cancer data reveals dosage-independent effects of differential expressed genes

To further demonstrate the ability of tICA to retrieve interesting patterns of variation in a multi-omic context, we applied it to the colon cancer TCGA dataset [1], comprising a matched subset of copy-number variation (CNV), DNA methylation and RNA-Seq data over 13971 genes and 272 samples (19 normals + 253 cancers) [29]. We applied tWFOBI to the resulting 3 × 272 × 13971 data-tensor, inferring a total of 3 × 37 ICs, which were ranked in order of decreasing kurtosis (**Methods**). Of the 37 ICs, 20 correlated with normal/cancer status (*P* < 0.05/37 ~ 0.001), with 4 of these capturing correlations between CNV and gene expression (**table.S1 in SI**). All 4 ICs were strongly enriched for specific chromosomal bands (**table.S1 in SI**), in line with those reported in the literature [1, 30], and one of these (IC-35) also exhibited concomitant correlation between DNAm and gene expression (**table.S1 in SI**). Plotting the weights of IC-35 along the CNV, DNAm and mRNA axes confirmed the ability of tWFOBI to identify patterns of mRNA expression variation which are driven by local CNV and which also associate with local variation in DNAm (**Fig.7A**). The corresponding weights along the sample-mode confirmed the association with normal-cancer status (**Fig.7B**), and scatterplots of the z-score normalized CNV and DNAm patterns against gene expression for one of the main driver genes (*STX6*) confirmed strong associations between CNV/DNAm and mRNA expression (**Fig.7C**). Strikingly, we observed that while variations in copy-number and DNAm of *STX6* modulate expression differences between colon cancers, that the deregulation of *STX6* expression between normal and cancer is clearly independent of copy-number and DNAm state (**Fig.7C**).

**Figure 7.**
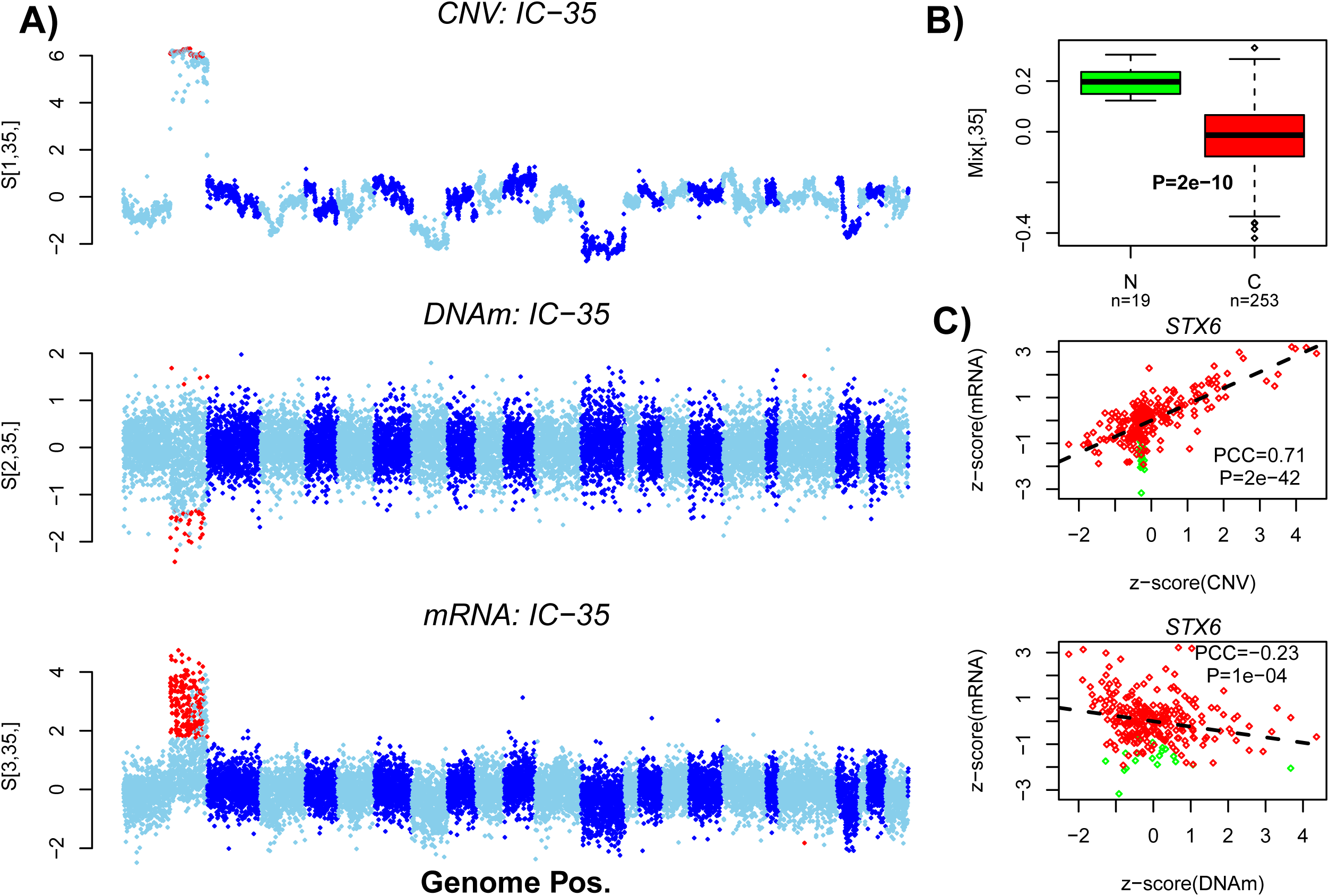
Validation of tensorial ICA on a multi-omic cancer set. **A)** Manhattan-like plots of IC-35 in gene-space, as inferred using tWFOBI on the colon TCGA set, projected along the CNV, DNAm and mRNA axes. Red points highlight genes that had large weights in both CNV and mRNA dimensions (CNV), in both DNAm and mRNA dimensions (DNAm), and the union of these (mRNA). Chromosomes are arranged in increasing order and displayed in alternating colors. **B)** Boxplots of the corresponding weights of IC-35 in sample-space, discriminating normal colon (N) from colon cancer (C). P-value is from a Wilcoxon rank sum test. **C)** Scatterplots of a driver gene (*STX6*) between z-score normalized segment level (CNV) and mRNA expression (top panel) and between z-score normalized DNAm level and mRNA expression (lower panel). Colors indicate normal (green) and cancer (red). Regression line, Pearson Correlation Coefficient and P-value are shown.

In order to validate this important finding and determine the extent of this phenomenon, we analysed 5 additional TCGA datasets (**Methods**), but now using a more direct approach: for each TCGA set, we first identified the subset of differentially expressed genes (DEGs) between normal and cancer (adjusted P-value threshold of 0.05), which also exhibit a positive correlation between expression and copy-number as assessed over cancers only, i.e. we selected those DEGs with a CNV-dosage effect across cancers. For those overexpressed in cancer, we then asked if individual tumours exhibiting a neutral CNV state (the CNV-state of the normal samples) or a CNV-loss, still exhibited overexpression relative to the normal samples. Remarkably, we observed that a very high fraction of these DEGs remained overexpressed when restricting to the subset of cancer samples with low or neutral CNV, thus indicating that *their overexpression in cancer is not dependent on CNV-state*, despite their expression across individual cancer samples being modulated by CNV-state (**Fig.8A**). This pattern of differential expression being independent of CNV-state was also seen in the case of DEGs with a CNV-dosage effect across tumours and which were underexpressed in cancer. Indeed, restricting to cancers with neutral or copy-number gain (**Fig.8A**), these genes were generally still underexpressed in these cancer samples compared to normal tissue. Similar patterns were observed when DEGs were selected for DNAm-expression dosage-effects across tumours (**Fig.8A**). Specific examples in the case of lung squamous cell carcinoma (LSCC) confirmed that DEGs in LSCC which exhibit a CNV or DNAm dosage effect across tumors, exhibit differential expression in a manner which is not dependent on CNV or DNAm state (**Fig.8B-C**). Thus, these data support the finding obtained using tICA, demonstrating the value and power of tICA to extract biologically important and novel patterns of data variation in a multi-omic context.

**Figure 8.**
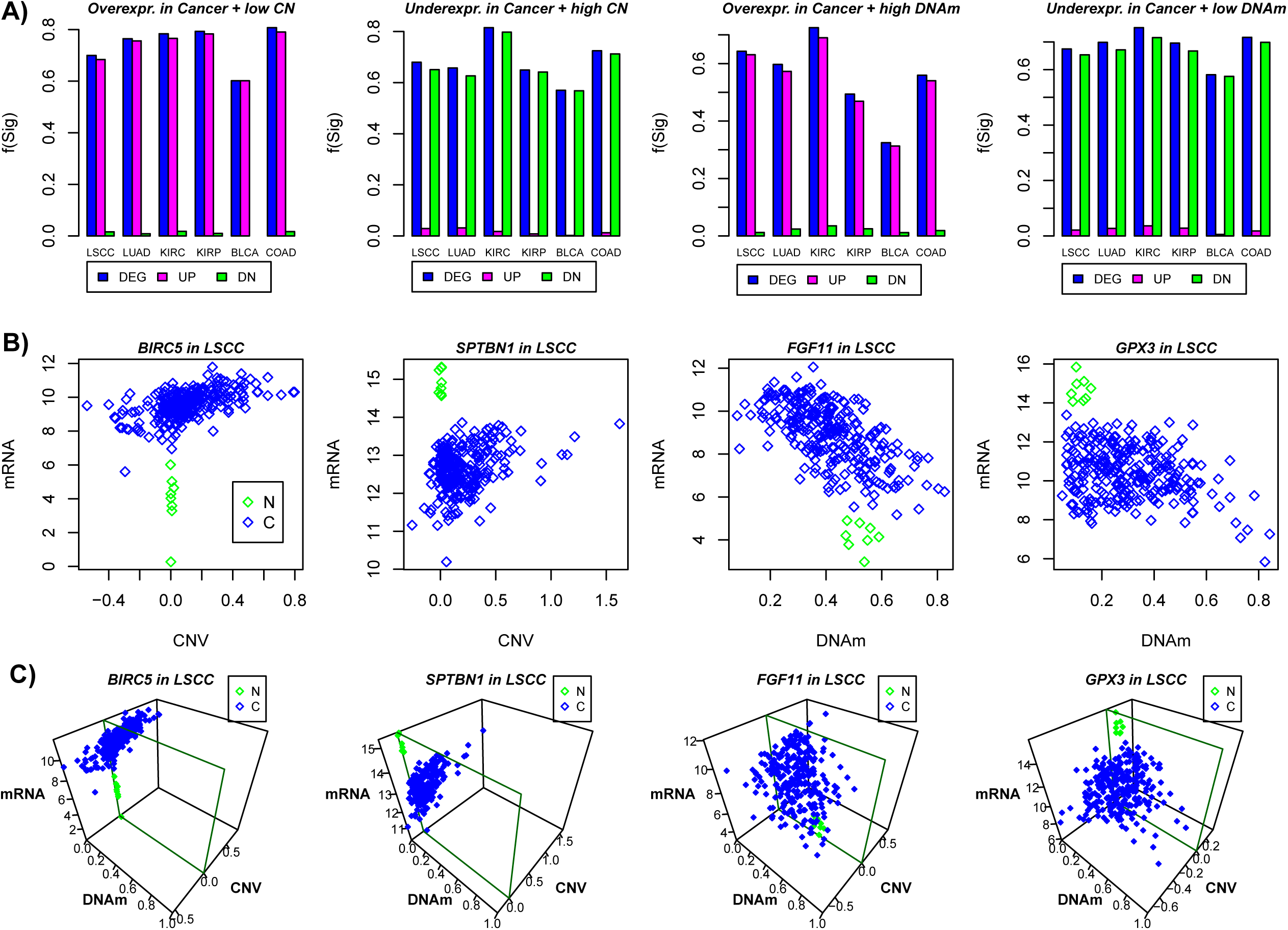
Multi-dimensional patterns of differential expression in cancer. **A)** Panels depict boxplots of the fraction of differentially expressed genes in cancer, which remain differentially expressed when specific cancer subsets are compared to normal-adjacent samples, for 6 different TCGA cancer types (LSCC, LUAD, KIRC, KIRP, BLCA, COAD), and for 4 different scenarios: genes overexpressed in cancer and considering cancers with neutral or copy-number loss of that gene (1st panel), genes underexpressed in cancer and considering cancers with neutral or copy-number gain (2nd panel), genes overexpressed in cancer and considering cancers with the highest levels of gene promoter DNAm (3rd panel), and finally genes underexpressed in cancer and considering cancers with the lowest levels of gene promoter DNAm (4th panel). In each panel, blue denotes the fraction of over/under expressed genes that are differentially expressed when only the specific cancer subset is compared to the normal samples, magenta denotes the fraction that are overexpressed whereas green denostes the fraction that are underexpressed. **B)** Scatterplots of mRNA expression against either copy-number variation level (CNV) or DNAm level for selected genes in LSCC. The selected genes represent examples of genes from A). For instance, BIRC5 in LSCC is overexpressed in cancer compared to normal, and this overexpression relative to normals is independent of the CNV of the cancer. **C)** As B), but now 3D scatterplots which also display the CNV or DNAm level. These plots illustrate that the difference in expression between cancer and normal is also independent of the other variable (e.g. CNV or DNAm). For instance, the underexpression of GPX3 in LSCC is neither driven by promoter DNAm, nor by CNV losses.

## Discussion

Here we have assessed and benchmarked a novel suite of tensorial decomposition algorithms (tPCA, tWFOBI, tWJADE) against a number of state-of-the-art alternatives. Specifically, while popular multi-way algorithms such as JIVE, iCluster or CCA/SCCA are in principle applicable to non-tensorial multi-way data (e.g. if the features across data-types are distinct or not matched), when assessed in a tensorial context (i.e when all dimensions are matched), these established methods are outperformed by the tensorial PCA and ICA methods considered here. This was demonstrated not only on simulated data, but also in the context of two real EWAS, where tICA methods were significantly more powerful to detect differentially methylated CpGs associated with an epidemiological factor (smoking) and SNPs (mQTLs). In the case of real EWAS, tICA also outperformed tPCA, in line with the fact that biological sources of data variation are non-Gaussian and sparse, and therefore more readily identified using statistical independence as a (non-linear) deconvolution criterion (as opposed to the linear decorrelation criterion used in tPCA). Thus, this extends the improvements seen for ICA over PCA on ordinary omic data matrices [13, 16] to the tensorial context. In addition, tPCA and tICA offer substantial (50–100 fold) speed advantages over methods like iCluster, JIVE and PARAFAC which can become computationally demanding or even prohibitive. Further application of tICA to a multi cell-type (B-cells, T-cells and monocytes) EWAS revealed its ability to identify loci enriched for cis-mQTLs (as cis-mQTLs make up over 90% of validated mQTLs in the ARIES database [25]). Indeed, tICA achieved relatively high sensitivity values with top-ranked CpGs in components containing over 60% mQTLs. Given that here we were limited by the fact that we did not have access to matched genotype information, our results demonstrate the potential of tICA to detect mQTLs in the absence of such genotype information. For instance, it identified many cell-type independent mQTLs, of which a substantial proportion validated in an independent blood-buccal EWAS study, and with several mapping to key GWAS loci for important diseases like Parkinson’s and schizophrenia. Although most of the identified mQTLs were blood cell-type independent, tICA estimated that approximately 25% of mQTLs may be blood cell-type specific, in line with the estimate of 20% obtained by Blueprint using a slightly different combination of blood cell subtypes (neutrophils, monocytes and T-cells) [28].

We note that application of tICA to any multi cell-type or multi-tissue EWAS is likely to have components strongly enriched for mQTLs, since for the same individuals DNAm is being measured in at least two different tissues or cell-types, and therefore genetic effects that do not depend on cell-type are bound to explain most of the inter-individual variation [31, 24]. Thus, we conclude that tICA could be an extremely versatile tool to identify novel candidate mQTLs in multi-cell EWAS for which matched genotype information may not be available. tICA may also help to identify groups of widely separated mQTLs which are regulated by the same SNP, and bound e.g. by a common transcription factor [32].

More generally, tICA can be applied to any multi-way data tensor to identify complex patterns of variation correlating with phenotypes of interest and the underlying features driving these variation patterns. This is accomplished by first correlating inferred independent components of variation in sample-mode space with sample phenotype information (e.g. age, smoking, normal/cancer status, genotype) and subsequently selecting the features with the largest weights in these correlated components. As an illustrative example, application of tICA to a multi-omic TCGA dataset revealed a deep novel insight: namely, that most differentially expressed genes in cancer *which exhibit a CNV or DNAm dosage-dependent effect on expression across individual tumors*, exhibit differential expression relative to the normal tissue in a manner which does not in fact depend on CNV or promoter DNAm state. In other words, although CNV and DNAm variation strongly modulates expression variation of these DEGs across individuals tumors, for most of the genes exhibiting this CNV or DNAm dosage-dependent expression pattern, their deregulation relative to normal cells appears to be independent of the underlying CNV or promoter DNAm state. Although it is clear that differential expression in cancer can be the result of many mechanisms other than CNV or DNAm, our observation is significant, because we did not just select cancer-DEGs, but the subset of these which exhibit a CNV or DNAm dosage-dependent effect on expression across tumours. The implications of our observation are important, given that many cancer classifications have been derived from unsupervised (clustering) analyses that were performed using only tumors, thus ignoring their patterns of variation relative to the normal reference state. Other large cancer studies, such as METABRIC [33], which did not profile normal tissue samples, identified candidate novel oncogenes and tumor suppressors solely on the basis of CNV-dosage effects on gene expression across cancers, yet our results indicate that this could identify many false positives in the sense that their overexpression or underexpression in cancer is not dependent on the underlying CNV-state. We point out that although this finding could have been obtained without application of a multi-way algorithm, that this would have required substantial prior insight. Therefore this subtle pattern of variation across multiple data-types was only discovered thanks to applying an agnostic method like tICA.

Although we have shown the value of tICA in identifying mQTLs and interesting patterns of variation across different data-types in cancer-genome data, it is also important to discuss some of the limitations, which however also apply to all other multi-way algorithms considered here. In particular, identifying sources of DNAm variation associated with epidemiological factors in a multi-tissue EWAS setting can be difficult due to confounding genetic variation. Indeed, in our application to a buccal-blood Illumina 450k EWAS we found that the sensitivity of all algorithms dropped very significantly if they are applied to all ~ 480,000 CpGs. Thus, it will be important in future to devise improvements to these tensorial methods. For instance, one solution may be to first perform dimensional reduction using supervised feature selection on separate data-types, and subsequently applying the tensorial methods on a reduced feature space. Alternatively, supervised tensorial methods such as tensorial Slice Inverse Regression [34] may help identify sources of variation specifically associated with epidemiological variables.

## Conclusions

In summary, the combined tPCA and tICA methods presented here will be an extremely valuable tool for analysis and interpretation of complex multi-way data, including multi-omic cancer data, as well as for the detection and clustering of mQTLs in multi cell-type EWAS where genotype information may not be available.

## Methods

Below we briefly describe the main tensorial BSS algorithms [8, 9, 35] as implemented here. For more technical details see [8, 9, 35]. We also provide brief details of our implementation of JIVE, PARAFAC, iCluster, CCA and sparse-CCA (SCCA). All these implementations are available as R-functions within **Additional File 2**.

### Tensorial PCA (tPCA)

We assume that we have *i* = 1, …, *p* i.i.d realizations of a matrix 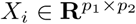, which can be structured as an order-3 data tensor *X* of dimension *p*_1_ × *p*_2_ × *p*. Then, tPCA decomposes *X* as follows:

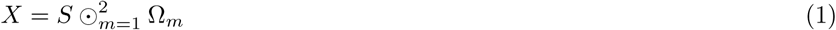

where *S* is also a 3-tensor of dimension *p*_1_ × *p*_2_ × *p*, Ω*_m_* (*m* = 1, 2) are orthogonal *p_m_* × *p_m_* matrices, i.e. 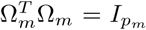, and where ☉ denotes the tensor contraction operator: for instance, for *Z* an *r*-tensor of dimension *p*_1_ × … × *p_r_* and *A* a matrix of dimension *p_m_* × *p_m_*, *Z* ☉*_m_ A* describes the *r*-tensor with entries 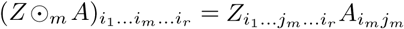 where the Einstein summation convention is assumed (i.e. indices appearing twice are summed over, e.g. *M_ik_M_in_* = *M_ik_M_in_* = (*M^T^ M*)*_kn_*). Thus, 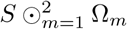 is a 3-tensor with entries

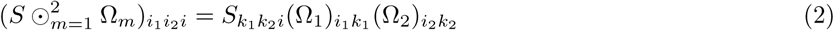

In the above tPCA decomposition, the entries 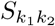 are assumed to be linearly uncorrelated. Introducing the operator ☉_−_*_m_*, which for general *r* is defined in entry form by

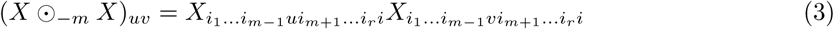

uncorrelated components means that the covariance matrix *S* ☉_−_*_m_ S* = Λ*_m_* is diagonal of dimension *p_m_* × *p_m_* and with entries the ranked eigenvalues of the *m*-mode covariance matrix (*X* ☉_−_*_m_ X*), which can be expressed as

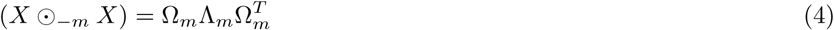

These ranked eigenvalues are useful for performing dimensional reduction, i.e. projecting the data onto subspaces carrying significant variation. For instance, one could use Random Matrix Theory (RMT) [36, 17] on each of the *m*-mode covariance matrices above to estimate the appropriate dimensionalities *d*_1_, …, *d_r_*. This would lead to a tPCA decomposition of the form 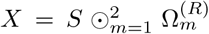, with *S* a *d*_1_ × *d*_2_ × *p* tensor and each 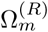 a reduced matrix obtained from Ω*_m_* by selecting the first *d_m_* columns. We note that for any of the original dimensions *p*_1_, … *p_r_* that are small, such dimensional reduction is not necessary.

In the applications considered here, our data tensor *X* is typically of dimension *n_t_* × *n_s_* × *n_G_*, where *n_t_* denotes the number of data or tissue types, *n_s_* the number of samples and *n_G_* the number of features (e.g. genes or CpGs). We note that the tPCA decomposition is performed on the first two dimensions (typically data-type and samples), so there are two relevant covariance matrices. In the special case of a data matrix (a 2-tensor), standard PCA involves the diagonalization of one data covariance matrix, hence for a 3-tensor there are two data covariance matrices, and for an (*r* + 1)-tensor, there are *r*. Here we use tPCA as implemented in the *tensorBSS* R-package [37].

### Tensorial ICA (tICA): the tWFOBI and tWJADE algorithms

For a data tensor 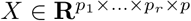 the tICA model is

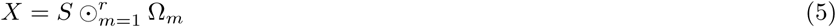

but now with the *p*_1_…*p_r_* random variables 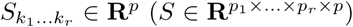 mutually statistically independent and satisfying 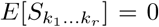 and 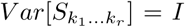. We note that *X* could be a suitably dimensionally reduced version *X*^(^*^R^*^)^ of *X*, such as one obtained using tPCA. For instance, in our applications, *X*^(^*^R^*^)^ would typically be a 3-tensor of dimension *n_t_* × *d_S_* × *n_G_* where *d_S_ < n_S_*. This dimensional reduction, and optionally the scaling of variances, is known as whitening (W).

As with ordinary ICA, there are different algorithms for inferring mutually statistical independent components 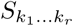. One algorithm is based on the concept of simultaneously maximising the fourth order moments (kurtosis) of the independent components (since by the Central Limit Theorem, linear mixtures of these are more Gaussian and therefore have smaller kurtosis values). This approach is known as fourth order blind identification (FOBI) [38]. Alternatively, one may attempt a joint approximate diagonalization of higher order eigenmatrices (JADE) [39, 35]. We note that although we use the tFOBI and tJADE functions in *tensorBSS*, that these do not implement tPCA beforehand. Hence, in this work we implement modified versions of tFOBI and tJADE which include a prior whitening transformation with tPCA. We call these modified versions tWFOBI & tWJADE.

### Benchmarking of tPCA and tICA against other tensor decomposition algorithms

JIVE (Joint and Individual Variation Explained) [5] is a powerful decomposition algorithm that identifies both joint and individual sources of data variation, i.e sources of variation that are common and specific to each data-type. For two data-types (i.e two tissue types or two types of molecular features), there are 3 key parameters that need to be specified or estimated in order to run JIVE: these are the number of components of joint variation (*dJ*) and the number of components of variation which are specific to each data-type (*dI*_1_, *dI*_2_). On simulated data, these parameters are chosen to be equal to the true (known) values, i.e. for our simulation model, *dJ* = 1, *dI*_1_=1 and *dI*_2_=1. In our real data applications, *dJ* is estimated using RMT on the concatanated matrix obtaining by merging the two data-type matrices together (after z-score normalising features in order to make them comparable), whilst *dI_i_* are estimated using RMT [17]. We note that these are likely upper bounds on the true number of individual sources of variation which are not also joint. We implemented JIVE using the *r.jive* R-package available from *http://www.r-project.org*.

PARAFAC (Parallel Factor Analysis) [6, 4] is a tensor decomposition algorithm whereby a data-tensor is decomposed into the sum of *R* terms, where each term is a factorised outer product of rank-1 tensors (i.e. vectors) over each mode. Thus, the one key parameter is *R* which equals the number of terms or components in the decomposition. In our simulation model, we chose *R* = 4. Although one of the two sources of variation in each data-type is common to both (hence 3 independent sources), we nevertheless ran PARAFAC with one additional component to more fairly assess its ability to infer components of joint variation. In the real data applications, we estimated *R* as 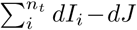 (with *n_t_* the number of tissue or cell-types), since this should approximately equal the total number of independent sources of variation. We implemented PARAFAC using the *multiway* R-package available from *http://www.r-project.org*.

iCluster [7] is a joint clustering algorithm for multi-way data, which models joint and individual sources of variation as latent Gaussian factors. The key parameter is K, which is the total number of clusters to infer. Although for the simulated data there were only 3 independent sources of variation, we chose *K* = 4 in order to more fairly assess the ability of the algorithm to infer the joint variation (choosing *K* = would “force” the algorithm to find the source of joint variation). We implemented iCluster using the *iCluster* R-package available from *http://www.r-project.org*.

Canonical Correlation Analysis (CCA) [20] and its sparse version, sparse-CCA (sCCA/SCCA) [21, 22], are methods to identify joint sources of variation (called canonical vectors) between two data matrices, where at least one of the dimensions is matched across data-types. Here we implement the version of CCA and sCCA of the R-package *PMA* available from *http://www.r-project.org*. One key parameter is *K*, the maximum number of canonical vectors to search for. Another parameter is the number of permutations used to estimate the significance of the covariance of each of the *K* canonical vectors. In each permutation, one of the data matrices is randomized (say by permuting the features around) and CCA/sCCA is reapplied. Since the data matrices are typically large, the distribution of covariances for the permuted cases is very tight, thus even 25 permutations is sufficient to estimate the number of significant canonical vectors reasonably well. The number of significant canonical vectors was defined as the number of components which exhibit observed covariances larger than the maximum value obtained over all 25 permutations, and is thus bounded above by *K*. In the non-sparse case, the two penalty parameters were chosen to be equal to 1, which means no penalty term is used. In the case of sCCA, we estimated best penalty parameters using an optimization procedure as described in [21, 22] with the number of permutations set to 25 and number of iterations equal to 15. On the simulated data, we ran CCA with *K* = 3, as *K* only needs to specify the maximum number of components to search for (the actual number of significant canonical vectors is 1 in our instance, as we have 1 source of joint variation). In the real data applications, we chose *K* to be equal to *dJ*, as estimated using the procedure for JIVE, and used a larger number of iterations (50) per run.

### Evaluation on simulated data

Here we describe the simulation model. The model first generates two data-matrices of dimension 1000 × 100, representing two data-types (e.g. DNA methylation and gene expression) where rows represent features and columns samples. We assume that the column and row labels (i.e. samples and genes) of the two matrices are identical and ordered in the same way. We assume one source of individual variation (IV) for each data matrix, each driven by 50 genes and 10 samples with the 50 genes and 10 samples unique to each data matrix. We also assume one source of common variation driven by a common set of 20 samples. The genes driving this common source of variation however, are assumed distinct for each data matrix. In total, there are 100 genes (50 for each data matrix) associated with this joint variation (JV). For the 50 genes driving the JV in one data-type and the 20 samples associated with this JV we draw the values from a Gaussian distribution 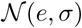, whereas for the other 50 genes in the other data-type we draw them from 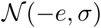, all with *e* = 3 and *σ* representing the noise level. Likewise for the IV we use Gaussians 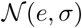. The rest of the data is modelled as noise 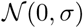. We consider a range of 9 noise levels, with *σ* ranging from 1 to 5 in steps of 0.5. Thus, at *σ* = 3, the *SNR* = *e*/*σ* = 1. For each noise level we perform 1000 Monte-Carlo runs, and for each run and algorithm we estimate the sensitivity (SE) and specificity (SP) for correctly identifying the 100 genes driving the JV. In the case of tPCA, tWJADE and tWFOBI, SE and SP were calculated as follows: we inferred a total of 12 components over the combined data-type and sample modes (2 in data-type mode × 6 in sample-space). We then projected the inferred components onto the original data-type dimensions, using the inferred 2 × 2 mixing matrix. For each data-type and each of the 6 components, we then selected the top-ranked 50 genes by absolute weight in the component. This allowed us to compute a SE and SP value for each data-type and component. For each component we then averaged the SE and SP values over the 2 data-types. In the last step, we select the component with the largest SE and SP value and record these values. We note that the resulting SE and SP values are not dependent on choosing 12 components. As long as the number of estimated components is larger than the total number of components of variation in the data (which for the simulated data is 4), results are invariant to the number of inferred components. In the case of CCA, which can only infer sources of joint variation, we ran it to infer a number of components (K=3) larger than the true one (only 1 source of JV). Pairs of canonical vectors were then selected according to whether their joint variance is larger than expected, as assessed using permutations. From hereon, the procedure to compute SE and SP proceeds as for the other algorithms, selecting the component with the best SE and SP value. As with the other methods, results do not depend on how we choose K as long as K is larger or equal than 1. In the case of PARAFAC, we ran it to infer R=4 components. Because for PARAFAC, there is only one inferred projection across features per component, for each component we rank the features according to their absolute weight, select the top-ranked 50, and then compute two separate SE (or SP) values, one for each of the 2 sets of 50 true positive genes driving JV. We then select for each set of JV driver genes, the component achieving the best SE (or SP). Finally, we average the SE and SP values for the two sets of true positives. As with the other algorithms, results do not depend on the choice of R, as long as R is larger or equal than 4 (since there 4 sources of variation: 2 of IV and 1 of JV which counts as 2 in the PARAFAC setting). In the case of JIVE, we ran it to infer 1 source of JV and 2 sources of IV. Because JIVE stacks the data matrices corresponding to the two data-types together, we then select the 100 top ranked genes, ranked by absolute weight in the inferred JV-matrix. SE and SP are then computed. Once again, results are stable to choosing a larger number of inferred sources of JV, because for the simulated data there is only 1 source of JV. Further details for all methods can be found in Additional File 2. Finally, for each algorithm and noise level, SE and SP are averaged over all the 1000 Monte-Carlo runs. Finally, statistical significance of SE and SP values between algorithms was assessed using paired non-parametric Wilcoxon rank sum tests. The whole analysis above was repeated for sources of variation drawn from a Laplace distribution (with same mean and standard deviation as the Gaussians above), in order to better capture the supergaussian nature of real biological data.

### Illumina 450k DNA methylation and multi-way TCGA datasets

We analysed Illumina 450k datasets from three main sources. One dataset is a multi blood cell subtype EWAS performed on 47 healthy individuals and 3 cell-types (B-cells, T-cells and Monocytes) [3]. Specifically, we used the same normalized data as used in [3], with the resulting data tensor being of dimension 3 × 47 × 388618, after removal of poor quality probes and probes with SNPs [40].

Another dataset was generated in [2], consisting of two tissue types (whole blood and buccal), 152 women and 447259 probes, resulting in a data tensor of dimension 2 × 152 × 447259. The 447259 probes is the number of probes obtained after quality control, removal of probes on X & Y chromosomes, polymorphic CpGs and probes with SNPs at the single-base extension site, and probes containing SNPs in their body as determined by Chen et al [40].

Finally, we also analysed 6 datasets from The Cancer Genome Atlas (TCGA). Specifically, we processed the RNA-Seq, Illumina 450k DNAm and copy-number data for 6 different cancer types (colon adenoma carcinoma (COAD), lung adenoma carcinoma (LUAD), lung squamous cell carinoma (LSCC), kidney renal cell carcinoma (KIRC), kidney papillary carcinoma (KIPC), bladder adenoma carcinoma (BLCA)), all of which contained reasonable number of normal-adjacent samples. The processing was carried out following the same procedure described by us in [29], which resulted in data-tensors over 3 data-types (mRNA, DNAm and copy-number), 14593 common genes and following sample numbers: 273 cancers and 8 normals for LSCC, 390 cancers and 20 normals for LUAD, 292 cancers and 24 normals for KIRC, 195 cancers and 21 normals for KIRP, 194 cancers and 13 normals for BLCA, and 253 cancers and 19 normals for COAD. We note that although these numbers of normal samples are small, that these are the normal samples with data for all 3 data-types.

### Identifying smoking-associated CpGs in the multi-tissue (whole blood + buccal) EWAS

In order to test the algorithms on real data we considered the matched multi-tissue (whole blood and buccal) Illumina 450k DNAm dataset for 152 women [2]. Smoking has been shown to be reproducibly associated with DNAm changes at a number of different loci [23]. We therefore used as a true positive set, a gold-standard list of 62 smoking-associated differentially methylated CpGs (smkDMCs), which have been shown to be correlated with smoking exposure in at least 3 independent whole blood EWAS [23]. The 62 smkCpGs were combined with 1000 randomly selected CpGs (non smoking-associated), resulting in a data-tensor of dimension 2 × 152 × 1062. Robustness was assessed by performing 1000 different Monte-Carlo runs, each run with a different random selection of 1000 non-smoking associated CpGs. The whole analysis was then repeated for 10000 randomly selected CpGs (data tensor of dimension 2 × 152 × 10062 and for a total of 1000 different Monte-Carlo runs. In the case of tPCA/tICA algorithms, the dimensionality parameters were chosen based on RMT as applied on the 2 separate matrices. Specifically, estimated unmixing matrices were of dimension 2×2 (for tissue-type mode) and *d×d* (for sample mode) with *d* the maximum of the two RMT estimates obtained from each tissue-type matrix. Sensitivity (SE) to capture the 62 smkCpGs was calculated in two different ways: in one approach we used the maximum SE attained by any IC, denoted SE(max), whilst in the other approach we allowed for the possibility that different enriched ICs could capture different subsets of smkCpGs. Thus, in the second approach, the SE was estimated by using the union of the selected CpGs over all enriched ICs. We note that enrichment of ICs for the smkCpGs was assessed using a simple binomial test and selecting those with a P-value less than the Bonferroni corrected value (ie. less than 0.05/number of ICs). In both approaches, the CpGs selected per component were the 62 with the largest absolute weights in the component, i.e. the number of selected CpGs was matched to the number of true positives.

In the case of JIVE, the number of components of joint variation was determined by applying RMT to the data matrix obtained by concatanating the features of the blood and buccal sets together with features standardised to unit variance to ensure comparability between data-types. For the number of components of individual variation we used the RMT estimates of each individual dataset, as this provides a safe upper bound. For PARAFAC, the number of components was determined by the sum of the RMT estimates for blood and buccal sets separately minus the value estimated for the concatanated matrix, as we reasoned that this would best approximate the total number of components of variation across the two data-types (joint or individual). For CCA and sCCA, the maximum number of canonical vectors to search for was set to be equal to the RMT estimate of the concatanated matrix, i.e. equal to the dimension of joint variation used in JIVE. For all methods, we selected the top-ranked 62 CpGs with largest absolute weights in each component, and estimated SE using the same two approaches described above for tPCA/tICA.

### mQTL and chromosome enrichment analysis

We applied tWFOBI to the data-tensor of a multi cell-type EWAS (Illumina 450k) performed on 47 healthy individuals and 3 cell-types (B-cells, T-cells and Monocytes) [3]. Specifically, we used the same normalized data as used in [3], i.e a data tensor of dimension 3×47×388618, after removal of poor quality probes and probes with SNPs [40]. Using RMT [17], we estimated a total of 11 components in sample mode-space, and so we inferred a source tensor of dimension 3 × 11, and mixing matrices of dimension 3 × 3 and 11 × 11. We also applied tWFOBI to the previous blood+buccal DNAm dataset, but for all 447259 probes that passed quality control. Applying RMT we estimated 26 significant components in sample-space. Hence, we applied tWFOBI on the 2 × 152 × 447259 data tensor to infer a source-tensor of dimension 2 × 26 × 447259 and mixing matrices of dimension 2 × 2 and 26 × 26. For both datasets, and for each inferred independent component, we selected the 500 probes with the largest absolute weights and tested enrichment of mQTLs against a high-confidence mQTL list (22245 mQTLs) from [25]. This list was generated as the overlap of mQTLs (passing a stringent P-value threshold of 1e-14) in blood derived from five different cohorts representing five different age groups. Odds ratios and P-values of enrichment were estimated using Fisher’s exact test. For chromosome enrichment, we obtained P-values using a binomial test. Concerning the selection of top-500 probes from each component, we note that this threshold is conservative, as all inferred ICs exhibited positive kurtosis with kurtosis values that remained significantly positive after removing the top-500 ranked probes.

To obtain estimates of cell-type independent and cell-type specific mQTLs, we used the following approach. The first mode/dimension of the estimated source tensor was rotated back to the original cell-types, using the estimated mixing matrix (of dimension 3 × 3, since there were 3 cell-types). For each of the previously enriched mQTLs, we compared its weights in all 3 components, each component being associated with a given cell type. For instance, if *S_t,cp,*_* denotes the component *cp* for cell-type t, thus defining a vector of weigths over all CpGs, we asked if the absolute weight of the given mQTL CpG is large for all cell-types or not. If sufficiently large (i.e. if within the top 10% quantile of the weight distribution) for all cell-types it was declared to be cell-type independent. If the mQTL weight for one or two cell-types fell within the lower 50% quantile of weights, we declared it a cell-type specific mQTL.

We also performed a comparative analysis of all multi-way algorithms in terms of their sensitivity to detect mQTLs, as given by the high-confidence list of 22245 mQTLs from the Aries database [25]. To assess the stability of the conclusions, we computed SE as described earlier, but considered a range of top number of selected CpGs per component, ranging from 500 up to 22245 in units of 500. As before, we estimated the overall SE taking into account the union of all selected CpGs from each components, as well as the maximum SE attained by any single component. Since the SE attained by any single component is bounded by the number of selected CpGs, we also considered the SE normalized for the number of selected CpGs.

### Application of tICA to multi-omic cancer data

We used the same normalized integrated copy-number state (segment values), Illumina 450k DNAm and RNA-Seq datasets of 6 cancer-types from the TCGA [1] and as used in our previous publication [29]. For the cancer-types considered see earlier subsection. We initially applied tWFOBI to the colon adenomacarcinoma TCGA dataset, estimating unmixing matrices of dimension 3 × 3 (for data-type) and *K ×K* (for sample mode) where *K* was the maximum RMT estimate over each of the 3 data-type matrices. Features driving each IC in each data-type dimension were selected using an iterative approach in which genes were ranked by absolute weight, and recursively removed until the kurtosis of the IC was less than 1, or the number of removed genes was larger than 500. Genes selected in common between the CNV and mRNA modes, or between the DNAm and mRNA modes, were declared “driver” genes between the respective data-types. To identify components correlating with normal-cancer status, we obtained the mixing matrix of the samples and then correlating each component to normal-cancer status using Wilcoxon’s rank sum test.

## Availability of data and materials

All data analysed here have already appeared in previous publications, or are publicly available. The Illumina 450k DNAm buccal-whole blood dataset from the NSHD published in [2] is available by submitting data requests to; see full policy at http://www.nshd.mrc.ac.uk/data.aspx. Managed access is in place for this 69 year old study to ensure that use of the data are within the bounds of consent given previously by participants, and to safeguard any potential threat to anonymity since the participants are all born in the same week [2]. The multi blood cell subtype Illumina 450k EWAS is available from the European Genome-phenome Archive (EGA) with the accession code EGAS00001001598 (https://www.ebi.ac.uk/ega/studies/EGAS00001001598) [3]. All TCGA data analysed here was downloaded and publicly available from the TCGA data portal website (https://portal.gdc.cancer.gov/). The ARIES mQTL database is available from (http://www.mqtldb.org) [25]. All software is available from the corresponding R-packages as described in Methods section, and their specific implementations as used in this manuscript, are available from github (https://github.com/jinghan1018/tensordecomp) and zenodo (https://zenodo.org/record/1208040 or https://doi.org/10.5281/zenodo.1208040) [41]. The tICA algorithms, as implemented here, are made available under a GNU General Public License version 3.

## Author’s contributions

AET devised the study, performed analyses and wrote manuscript. JH performed statistical analyses. JV and KN contributed methodology and to the writing of the manuscript. DSP contributed and prepared EWAS datasets for statistical analyses

## Ethics approval

No ethics approval was necessary for this publication as all data analyzed is already freely available in the public domain.

## References

1. TCGA: Comprehensive molecular characterization of human colon and rectal cancer. Nature 487(7407), 330–7 (2012)

2. Teschendorff, A.E., Yang, Z., Wong, A., Pipinikas, C.P., Jiao, Y., Jones, A., Anjum, S., Hardy, R., Salvesen, H.B., Thirlwell, C., Janes, S.M., Kuh, D., Widschwendter, M.: Correlation of smoking-associated dna methylation changes in buccal cells with dna methylation changes in epithelial cancer. JAMA Oncol 1(4), 476–85 (2015)

3. Paul, D.S., Teschendorff, A.E., Dang, M.A., Lowe, R., Hawa, M.I., Ecker, S., Beyan, H., Cunningham, S., Fouts, A.R., Ramelius, A., Burden, F., Farrow, S., Rowlston, S., Rehnstrom, K., Frontini, M., Downes, K., Busche, S., Cheung, W.A., Ge, B., Simon, M.M., Bujold, D., Kwan, T., Bourque, G., Datta, A., Lowy, E., Clarke, L., Flicek, P., Libertini, E., Heath, S., Gut, M., Gut, I.G., Ouwehand, W.H., Pastinen, T., Soranzo, N., Hofer, S.E., Karges, B., Meissner, T., Boehm, B.O., Cilio, C., Larsson, H.E., Lernmark, A., Steck, A.K., Rakyan, V.K., Beck, S., Leslie, R.D.: Increased dna methylation variability in type 1 diabetes across three immune effector cell types. Nat Commun 7, 13555 (2016)

4. Hore, V., Vinuela, A., Buil, A., Knight, J., McCarthy, M.I., Small, K., Marchini, J.: Tensor decomposition for multiple-tissue gene expression experiments. Nat Genet 48(9), 1094–100 (2016)

5. Lock, E.F., Hoadley, K.A., Marron, J.S., Nobel, A.B.: Joint and individual variation explained (jive) for integrated analysis of multiple data types. Ann Appl Stat 7(1), 523–542 (2013)

6. Bro, R.: Parafac. tutorial and applications. Chem Intel Lab Syst 38, 149–171 (1997)

7. Shen, R., Olshen, A.B., Ladanyi, M.: Integrative clustering of multiple genomic data types using a joint latent variable model with application to breast and lung cancer subtype analysis. Bioinformatics 25(22), 2906–12 (2009)

8. Virta, J., Taskinen, S., Nordhausen, K.: Applying fully tensorial ica to fmri data. In: Signal Processing in Medicine and Biology Symposium (SPMB), 2016 IEEE, pp. 1–6 (2016). IEEE

9. Virta, J., Li, B., Nordhausen, K., Oja, H.: Independent component analysis for tensor-valued data. Journal of Multivariate Analysis 162, 172–192 (2017)

10. Comon, P.: Independent component analysis, a new concept? Signal Process. 36(3), 287–314 (1994)

11. Liebermeister, W.: Linear modes of gene expression determined by independent component analysis. Bioinformatics 18(1), 51–60 (2002)

12. Martoglio, A.M., Miskin, J.W., Smith, S.K., MacKay, D.J.: A decomposition model to track gene expression signatures: preview on observer-independent classification of ovarian cancer. Bioinformatics 18(12), 1617–1624 (2002)

13. Teschendorff, A.E., Journ´ee, M., Absil, P.A., Sepulchre, R., Caldas, C.: Elucidating the altered transcriptional programs in breast cancer using independent component analysis. PLoS Comput Biol 3(8), 161 (2007)

14. Kowarsch, A., Blochl, F., Bohl, S., Saile, M., Gretz, N., Klingmuller, U., Theis, F.J.: Knowledge-based matrix factorization temporally resolves the cellular responses to il-6 stimulation. BMC Bioinformatics 11, 585 (2010)

15. Illner, K., Fuchs, C., Theis, F.J.: Bayesian blind source separation for data with network structure. J Comput Biol 21(11), 855–65 (2014)

16. Biton, A., Bernard-Pierrot, I., Lou, Y., Krucker, C., Chapeaublanc, E., Rubio-Perez, C., Lopez-Bigas, N., Kamoun, A., Neuzillet, Y., Gestraud, P., Grieco, L., Rebouissou, S., de Reynies, A., Benhamou, S., Lebret, T., Southgate, J., Barillot, E., Allory, Y., Zinovyev, A., Radvanyi, F.: Independent component analysis uncovers the landscape of the bladder tumor transcriptome and reveals insights into luminal and basal subtypes. Cell Rep 9(4), 1235–45 (2014)

17. Teschendorff, A.E., Zhuang, J., Widschwendter, M.: Independent surrogate variable analysis to deconvolve confounding factors in large-scale microarray profiling studies. Bioinformatics 27(11), 1496–1505 (2011)

18. Alexandrov, L.B., Nik-Zainal, S., Wedge, D.C., Campbell, P.J., Stratton, M.R.: Deciphering signatures of mutational processes operative in human cancer. Cell Rep 3(1), 246–259 (2013)

19. Zhang, S., Liu, C.C., Li, W., Shen, H., Laird, P.W., Zhou, X.J.: Discovery of multi-dimensional modules by integrative analysis of cancer genomic data. Nucleic Acids Res 40(19), 9379–9391 (2012)

20. Hotelling, H.: Relations between two sets of variates. Biometrika 28(3–4), 321–377 (1936)

21. Witten, D.M., Tibshirani, R.J.: Extensions of sparse canonical correlation analysis with applications to genomic data. Stat Appl Genet Mol Biol 8, 28 (2009)

22. Witten, D.M., Tibshirani, R., Hastie, T.: A penalized matrix decomposition, with applications to sparse principal components and canonical correlation analysis. Biostatistics 10(3), 515–34 (2009)

23. Gao, X., Jia, M., Zhang, Y., Breitling, L.P., Brenner, H.: Dna methylation changes of whole blood cells in response to active smoking exposure in adults: a systematic review of dna methylation studies. Clin Epigenetics 7, 113 (2015)

24. van Dongen, J., Nivard, M.G., Willemsen, G., Hottenga, J.J., Helmer, Q., Dolan, C.V., Ehli, E.A., Davies, G.E., van Iterson, M., Breeze, C.E., Beck, S., Suchiman, H.E., Jansen, R., van Meurs, J.B., Heijmans, B.T., Slagboom, P.E., Boomsma, D.I.: Genetic and environmental influences interact with age and sex in shaping the human methylome. Nat Commun 7, 11115 (2016)

25. Gaunt, T.R., Shihab, H.A., Hemani, G., Min, J.L., Woodward, G., Lyttleton, O., Zheng, J., Duggirala, A., McArdle, W.L., Ho, K., Ring, S.M., Evans, D.M., Smith, G.D., Relton, C.L.: Systematic identification of genetic influences on methylation across the human life course. Genome Biol 17, 61 (2016)

26. Satake, W., Nakabayashi, Y., Mizuta, I., Hirota, Y., Ito, C., Kubo, M., Kawaguchi, T., Tsunoda, T., Watanabe, M., Takeda, A., Tomiyama, H., Nakashima, K., Hasegawa, K., Obata, F., Yoshikawa, T., Kawakami, H., Sakoda, S., Yamamoto, M., Hattori, N., Murata, M., Nakamura, Y., Toda, T.: Genome-wide association study identifies common variants at four loci as genetic risk factors for parkinson’s disease. Nat Genet 41(12), 1303–7 (2009)

27. Kahler, A.K., Djurovic, S., Kulle, B., Jonsson, E.G., Agartz, I., Hall, H., Opjordsmoen, S., Jakobsen, K.D., Hansen, T., Melle, I., Werge, T., Steen, V.M., Andreassen, O.A.: Association analysis of schizophrenia on 18 genes involved in neuronal migration: Mdga1 as a new susceptibility gene. Am J Med Genet B Neuropsychiatr Genet (7), 1089–100 (2008)

28. Chen, L., Ge, B., Casale, F.P., Vasquez, L., Kwan, T., Garrido-Martin, D., Watt, S., Yan, Y., Kundu, K., Ecker, S., Datta, A., Richardson, D., Burden, F., Mead, D., Mann, A.L., Fernandez, J.M., Rowlston, S., Wilder, S.P., Farrow, S., Shao, X., Lambourne, J.J., Redensek, A., Albers, C.A., Amstislavskiy, V., Ashford, S., Berentsen, K., Bomba, L., Bourque, G., Bujold, D., Busche, S., Caron, M., Chen, S.H., Cheung, W., Delaneau, O., Dermitzakis, E.T., Elding, H., Colgiu, I., Bagger, F.O., Flicek, P., Habibi, E., Iotchkova, V., Janssen-Megens, E., Kim, B., Lehrach, H., Lowy, E., Mandoli, A., Matarese, F., Maurano, M.T., Morris, J.A., Pancaldi, V., Pourfarzad, F., Rehnstrom, K., Rendon, A., Risch, T., Sharifi, N., Simon, M.M., Sultan, M., Valencia, A., Walter, K., Wang, S.Y., Frontini, M., Antonarakis, S.E., Clarke, L., Yaspo, M.L., Beck, S., Guigo, R., Rico, D., Martens, J.H., Ouwehand, W.H., Kuijpers, T.W., Paul, D.S., Stunnenberg, H.G., Stegle, O., Downes, K., Pastinen, T., Soranzo, N.: Genetic drivers of epigenetic and transcriptional variation in human immune cells. Cell 167(5), 1398–1414 (2016)

29. Teschendorff, A.E., Zheng, S.C., Feber, A., Yang, Z., Beck, S., Widschwendter, M.: The multi-omic landscape of transcription factor inactivation in cancer. Genome Med 8(1), 89 (2016)

30. Mayrhofer, M., Kultima, H.G., Birgisson, H., Sundstrom, M., Mathot, L., Edlund, K., Viklund, B., Sjoblom, T., Botling, J., Micke, P., Pahlman, L., Glimelius, B., Isaksson, A.: 1p36 deletion is a marker for tumour dissemination in microsatellite stable stage ii-iii colon cancer. BMC Cancer 14, 872 (2014)

31. Teschendorff, A.E., Relton, C.L.: Statistical and integrative system-level analysis of dna methylation data. Nat Rev Genet 19(3), 129–147 (2018)

32. Bonder, M.J., Luijk, R., Zhernakova, D.V., Moed, M., Deelen, P., Vermaat, M., van Iterson, M., van Dijk, F., van Galen, M., Bot, J., Slieker, R.C., Jhamai, P.M., Verbiest, M., Suchiman, H.E., Verkerk, M., van der Breggen, R., van Rooij, J., Lakenberg, N., Arindrarto, W., Kielbasa, S.M., Jonkers, I., van’t Hoof, P., Nooren, I., Beekman, M., Deelen, J., van Heemst, D., Zhernakova, A., Tigchelaar, E.F., Swertz, M.A., Hofman, A., Uitterlinden, A.G., Pool, R., van Dongen, J., Hottenga, J.J., Stehouwer, C.D., van der Kallen, C.J., Schalkwijk, C.G., van den Berg, L.H., van Zwet, E.W., Mei, H., Li, Y., Lemire, M., Hudson, T.J., Slagboom, P.E., Wijmenga, C., Veldink, J.H., van Greevenbroek, M.M., van Duijn, C.M., Boomsma, D.I., Isaacs, A., Jansen, R., van Meurs, J.B., van’t Hoen, P.A., Franke, L., Heijmans, B.T.: Disease variants alter transcription factor levels and methylation of their binding sites. Nat Genet 49(1), 131–138 (2017)

33. Curtis, C., Shah, S.P., Chin, S.F., Turashvili, G., Rueda, O.M., Dunning, M.J., Speed, D., Lynch, A.G., Samarajiwa, S., Yuan, Y., Gr¨af, S., Ha, G., Haffari, G., Bashashati, A., Russell, R., McKinney, S., A, A.L., A, A.G., Provenzano, E., G, G.W., Pinder, S., Watson, P., Markowetz, F., Murphy, L., Ellis, I., Purushotham, A., Borresen-Dale, A.L., Brenton, J.D., Tavare, S., Caldas, C., Aparicio, S.: The genomic and transcriptomic architecture of 2,000 breast tumours reveals novel subgroups. Nature 486(7403), 346–352 (2012)

34. Ding, S., Cook, R.D.: Tensor sliced inverse regression. J. Multivariate Analysis 133, 216–231 (2015)

35. Virta, J., Li, B., Nordhausen, K., Oja, H.: JADE for tensor-valued observations. Accepted to Journal of Computational and Graphical Statistics (2017). doi:10.1080/10618600.2017.1407324. arXiv preprint arXiv:1603.05406

36. Plerou, V., Gopikrishnan, P., Rosenow, B., Amaral, L.A., Guhr, T., Stanley, H.E.: Random matrix approach to cross correlations in financial data. Phys Rev E Stat Nonlin Soft Matter Phys 65(6), 066126 (2002)

37. Virta, J., Li, B., Nordhausen, K., Oja, H.: tensorBSS: Blind Source Separation Methods for Tensor-Valued Observations. (2017). R package version 0.3. https://CRAN.R-project.org/package=tensorBSS

38. Cardoso, J.-F.: Source separation using higher order moments. In: Acoustics, Speech, and Signal Processing, 1989. ICASSP-89., 1989 International Conference On, pp. 2109–2112 (1989). IEEE

39. Cardoso, J.F., Souloumiac, A.: Blind beamforming for non gaussian signals. IEE Proceedings-F 140, 362–370 (1993)

40. Chen, Y.A., Lemire, M., Choufani, S., Butcher, D.T., Grafodatskaya, D., Zanke, B.W., Gallinger, S., Hudson, T.J., Weksberg, R.: Discovery of cross-reactive probes and polymorphic cpgs in the illumina infinium humanmethylation450 microarray. Epigenetics 8(2), 203–9 (2013)

41. Jing, H., Teschendorff, E.A.: R-scripts for implementing tensor decomposition methods (2018). doi:10.5281/zenodo.1208040. https://doi.org/10.5281/zenodo.1208040

